# Genome-wide association study and genetic differentiation analysis reveal the genetic basis of *Vibrio harveyi* resistance in blackspotted croaker (*Protonibea diacanthus*)

**DOI:** 10.64898/2026.09.10.750604

**Authors:** Song Sun, Liang Zhong, Lin Yan, Kuoqiu Yan, Ning Li, Peng Xu, Wenlong Cai

## Abstract

*Vibrio harveyi* is a primary causative agent of vibriosis, leading to catastrophic economic losses in the blackspotted croaker (*Protonibea diacanthus*) aquaculture. However, the genetic architecture underlying its resistance remains poorly understood. In this study, we conducted the first integrated genome-wide association study (GWAS) and genetic differentiation analysis (*F*_ST_) to investigate the genetic basis of *V. harveyi* resistance. A total of 450 juveniles were challenged, and 300 individuals (comprising 150 resistant and 150 susceptible) were selected for whole-genome resequencing. Genomic heritability (*h*^*2*^) was estimated at 0.374±0.29 for survival time (ST) and 0.237±0.185 for survival status (SS), indicating moderate genetic control. Notably, an exceptionally high genetic correlation (*r*_*g*_ = 0.999) was observed between ST and SS, while resistance traits were genetically independent of growth performance (*r*_*g*_ < 0.03). Through the convergence of GWAS and selection sweep signals, a major-effect QTL was identified on chromosome 4. Candidate gene prioritization identified high-confidence targets involved in post-transcriptional regulation and mRNA stability, specifically *elavl4* (harboring a top-tier exon variant), *rorb, cbpc6*, and *epn4*. Functional enrichment analysis further revealed that resistance is significantly associated with “rRNA processing”, “mRNA surveillance pathway”, and “nucleocytoplasmic transport”, suggesting that efficient RNA metabolic control is a critical determinant of host survival. These findings provide novel insights into the non-canonical immune mechanisms of *P. diacanthus* and offer robust molecular markers for marker-assisted selection and genomic selection, facilitating the development of disease-resistant strains in the sciaenid industry.

## 1. Introduction

The blackspotted croaker (*Protonibea diacanthus*), belonging to the Sciaenidae, is a valuable fish species found along the southeastern coast of China, and its aquaculture industry is mainly distributed in coastal areas such as Guangdong and Fujian. As a high-value sciaenid, *P. diacanthus* has become a prime target for large-scale aquaculture, largely owing to its desirable nutritional profile and the lucrative returns associated with its gas bladder. In recent years, the continuous decline of wild stocks coupled with surging market demand has spurred significant interest in their artificial propagation and large-scale farming. However, under high-density intensive aquaculture conditions, the industry faces severe challenges from infectious diseases, which have become a primary bottleneck restricting its sustainable development.

Among the various pathogens affecting mariculture, *Vibrio harveyi* stands out as one of the most destructive, capable of infecting a diverse array of teleosts and crustaceans. It is the causative agent of vibriosis, characterized by acute septicemia and mass mortality. Outbreaks have been documented globally across numerous farmed species, including the yellow drum (*Nibea albiflora*) (Luo et al., 2021), Chinese tongue sole (*Cynoglossus semilaevis*) (Li et al., 2019), leopard coral grouper (*Plectropomus leopardus*) (Lu et al., 2023), golden pompano (*Trachinotus ovatus*) (Cao et al., 2018), and the lined seahorse (*Hippocampus erectus*) (Li et al., 2024). In *C. semilaevis* aquaculture, *V. harveyi* infections can result in devastating mortality rates of 50–70% (Li et al., 2019). As documented by Deng et al. (2019), *V. harveyi* exhibits substantial genomic plasticity, enabling the uptake of diverse virulence factors and resistance genes through horizontal flow, which ultimately drives its adaptive evolution and virulence. The marked heterogeneity in serological and biochemical characteristics among strains from different geographical origins further complicates disease management (Pavlinec et al., 2022).

Conventional strategies for mitigating vibriosis primarily rely on antibiotic treatment, vaccination, and environmental management. While antibiotics can offer short-term control, their excessive use has led to widespread antimicrobial resistance. Strains isolated from shrimp aquaculture environments have demonstrated broad resistance to multiple antibiotics, posing significant risks to food safety and ecological integrity (Stalin & Srinivasan, 2016). Regarding immunoprophylaxis, the efficacy of vaccines remains limited by the high serological diversity of *Vibrio* species and the absence of a conventional adaptive immune system in certain aquaculture taxa, such as mollusks (Yang et al., 2024). Consequently, enhancing the innate disease resistance of the host through genetic improvement is increasingly recognized as the most sustainable and fundamental approach for long-term disease control.

To achieve long-term improvements in disease resistance within aquaculture, genetic selection represents a pivotal approach. By evaluating single nucleotide polymorphism (SNP) variations across the whole genome, genome-wide association studies (GWAS) enable researchers to unravel the complex genetic underpinnings of these defensive traits. This approach facilitates the identification of significant loci and candidate genes, providing crucial molecular targets for marker-assisted selection (MAS) and genomic selection (GS). In recent years, the implementation of GWAS-based selection strategies has markedly advanced the development of aquaculture lineages resistant to *V. harveyi* infection. In the yellow drum, a closely related sciaenid, GWAS identified 23 significant SNPs and 8 candidate genes (including *sphk, cacna1h*, and *mgat5b*) on chromosome 14, explaining 10.64%–13.68% of the phenotypic variance (Luo et al., 2021). Building on this, integrative GWAS and eQTL analysis identified *socs1, clec16a, ciita*, and *prkcb* as key immune-related candidates (Huang et al., 2024). Subsequent functional validations have elucidated the roles of several candidate genes in the immune response against *V. harveyi*, including antimicrobial peptides like *NK-lysin* (Huang et al., 2023), *Galectin-9* (Luo et al., 2023), *sphk1* and its interaction with *TFAP2A* (Cui et al., 2024), *Cytoglobin* (*Cygb*) (Zhou et al., 2023), *G-type lysozyme* (Xiao et al., 2024), and the *perforin-1 like* gene (Fu et al., 2021). In *C. semilaevis*, genetic parameters for *V. harveyi* resistance were estimated through challenge tests (Li et al., 2019), and the feasibility of incorporating resistance traits into breeding programs was validated using natural outbreak data (Hu et al., 2020). Integrative genomics and transcriptomics have further revealed novel molecular mechanisms of vibriosis resistance in this species (Zhou et al., 2022). Analogous genomic strategies have been successfully implemented across diverse aquatic models; for instance, genome-wide association testing and predictive modeling were utilized to investigate enteritis susceptibility in the lined seahorse (Li et al., 2024) and survival performance in the leopard coral grouper (Lu et al., 2023). These studies collectively demonstrate that resistance to *V. harveyi* is a heritable trait, and GWAS provides a viable technical pathway for molecular breeding. Despite the wealth of data on related sciaenids like *Larimichthys crocea* (Zhao et al., 2021), the genetic basis of disease resistance in *P. diacanthus* remains entirely unexplored. Given the differences in genetic background, immune system composition, and host-pathogen interactions across species, findings from other fish cannot be directly extrapolated to the blackspotted croaker. Therefore, there is an urgent need to establish a species-specific genetic analysis framework for this high-value fish.

In the present study, we conducted a controlled *V. harveyi* challenge in a population of *P. diacanthus* and performed a GWAS using 150 resistant and 150 susceptible individuals. The core goals of this investigation were threefold: first, to quantify the genomic heritability governing vibriosis resistance phenotypes in *P. diacanthus*; second, to pinpoint prominent genomic intervals along with high-confidence candidate genes linked to survival outcomes; and third, to lay down a vital molecular groundwork for subsequent marker-aided and genomic selection frameworks. Ultimately, our discoveries are intended to bridge the prevailing information void in *P. diacanthus* genetics while offering valuable perspectives for breeding disease-resistant strains across the Sciaenidae family.

## 2. Materials and methods

### 2.1. Sample collection and *Vibrio harveyi* challenge

Juvenile blackspotted croaker (*Protonibea diacanthus*) were obtained from Guangdong Bluegen Marine Biotechnology Company. Before the experiment, six fish were randomly sampled for a comprehensive health assessment. The fish were confirmed to be pathogen-free, as evidenced by the absence of parasites, no bacterial growth from internal organs, and negative PCR results for common viruses (*GIV, VNNV*, and *ISKNV*). Prior to the trial, the fish underwent a 14-day adaptation period inside 1000 L fiberglass-reinforced plastic (FRP) containers. Throughout this conditioning phase, the cohort was supplied with feed twice a day (specifically at 09:00 and 17:00) until visual satiety was reached. To ensure excellent system hygiene, a complete (100%) water replacement was executed on a daily basis. The physicochemical variables were strictly regulated within the following thresholds: temperature at 29.21 ± 0.22□, salinity 27.05 ± 0.3‰, pH 7.5-8.0, and dissolved oxygen > 6.0 mg/L. Ammonia-N and nitrite levels were kept below 0.1 mg/L.

A virulent strain of *V. harveyi*, isolated from diseased *P. diacanthus*, was used for the challenge. To ensure optimal virulence, the strain was revived on TCBS agar at 28 □ for 24 h. Subsequently, an isolated colony was transferred into Luria-Bertani (LB) medium and incubated at 28 °C for a duration of 16-20 h. Biomass collection was achieved via centrifugation at 4,000 rpm for 10 min, after which the bacterial pellet was washed and mixed into sterile PBS. Utilizing benchmarks from our prior LD_50_. evaluations, the final bacterial titer was adjusted to a working concentration of 3.2 × 10^8^ CFU/mL.

A total of 450 healthy 100-day-old juveniles (mean length 12.56±2.01 cm; mean weight 37.8±15.36 g) were randomly distributed into five 1000 L FRP tanks (90 fish per tank). After 24 h of fasting, each fish received an intraperitoneal (i.p.) injection of 100 μL of the bacterial suspension. Concurrently, a negative control cohort (n = 45) was administered an equivalent volume of sterile PBS. Following injection, mortality and clinical signs were monitored every 2-3 h. Dead individuals (defined as the susceptible group) were recorded and sampled immediately. The monitoring ceased when no mortality occurred for three consecutive days, by which time 240 dead fish had been collected. The remaining survivors (defined as the resistant group) were maintained for an additional three weeks. Fin clips from all individuals were preserved in absolute ethanol for subsequent genomic DNA extraction.

During the bacterial challenge, each fish was individually identified and monitored. At the time of sampling or recording mortality, the standard length (SL) and body weight (BW) of each individual were measured and recorded. The survival status (1 for survived, 0 for dead) and the specific survival time (measured in hours post-infection, hpi) were documented for every fish. The experiment was officially terminated at 444 h, and all individuals that survived until the end of the trial were assigned a maximum survival time of 444 h as right-censored data. A total of 300 individuals were selected for whole-genome resequencing (WGR) using an extreme phenotype sampling strategy. This sequencing panel consisted of 150 susceptible individuals, randomly chosen from the 240 fish that died during the acute challenge, and 150 resistant individuals, randomly selected from the survivors that remained healthy for over three weeks. The comprehensive individual profiles—comprising morphometric traits (SL and BW) and survival parameters (status and time)—were strictly linked to their respective genomic data to facilitate statistical correction for growth-related effects in subsequent association analyses. The animal study was approved and conducted according to the guidelines of the Animal Experiment Committee of City University of Hong Kong.

### 2.2. Whole-genome Resequencing and Variant Calling

Total genomic DNA isolation was performed utilizing a commercial TIANamp Marine Animals DNA Kit (Tiangen, China). The concentration, structural intactness, and chemical purity of each sample were subsequently verified via fluorometric measurements alongside 1% agarose gel electrophoresis (run at 150 V for 40 min). To initiate sequencing library construction, input DNA quantities ranging from 300 ng to 1 µg were sheared via enzymatic cleavage. The resulting fragments were size-selected using magnetic beads to enrich for DNA in the 200-400 bp range. The sequencing library was constructed using the Hieff NGS® OnePot Pro DNA Library Prep Kit V4 (12972). DNA fragments underwent end repair, A-tailing, and adapter ligation to the 3’ ends under controlled temperature conditions. The adapter-ligated products were amplified via PCR and purified using a magnetic bead-based cleanup. To generate the final library, PCR products were denatured into single strands and circularized under optimized conditions. Uncircularized linear DNA molecules were subsequently digested to yield a single-stranded circular library. Qualified libraries were sequenced on the DNBSEQ platform. Single-stranded circular DNA molecules were amplified via rolling circle replication (RCR) to form DNA nanoballs (DNBs), each containing more than 300 copies. These DNBs were loaded into nanowells on a high-density DNA nanochip and sequenced using combinatorial Probe-Anchor Synthesis (CPAS) technology.

Raw reads were filtered via Trimmomatic and aligned to the *P. diacanthus* reference genome (GCA_028641955.1) (Xu et al., 2022) using BWA. Variant calling was performed with the GATK (v4.6.1.0) pipeline. To ensure high-quality markers, variants were subjected to strict hard filtering (QD<2.0, MQ < 40.0, FS>60.0, SOR>3.0, MQRankSum<-12.5, and ReadPosRankSum < −8.0). This process yielded 3,743,916 high-quality SNPs and 602,790 InDels with an 86% retention rate, providing a robust dataset for subsequent association mapping. Post-call filtering of SNPs was conducted using PLINK (v1.9) (Purcell et al., 2007). Variant filtering was performed using PLINK (v1.9) and VCFtools. Criteria included: minor allele frequency (MAF) > 0.05, maximum missing rate per site < 0.1, and Hardy-Weinberg equilibrium (HWE) P > 1×10^-6^.

### 2.3. Population structure and genetic relatedness analysis

To account for potential confounding effects in the association mapping, population stratification was evaluated via principal component analysis (PCA) using PLINK (v1.90b6.21). The analysis was performed using the pruned set of high-quality SNPs to minimize the impact of linkage disequilibrium. The PCA results were visualized as a scatter plot of the first two principal components (PC1 vs. PC2) using the ggplot2 package (v3.5.2) in R to assess the clustering patterns and ensure population homogeneity. Additionally, genetic relatedness among the 300 individuals was quantified by constructing a genomic relationship matrix (GRM). The GRM was calculated using GEMMA (v0.98.5) with the -gk 1 parameter, which accounts for pairwise genetic similarity based on the centralized genotype matrix. To illustrate the kinship and genetic proximity within the experimental population, the GRM was visualized as a heatmap using the pheatmap package in R. This integrated approach ensured that any observed phenotypic variations were linked to genetic resistance rather than cryptic relatedness or population structure.

### 2.4. Genome-wide Association Study (GWAS)

Genome-wide association studies (GWAS) were performed using the Fixed and Random Model Circulating Probability Unification (FarmCPU) method (Liu et al., 2022), implemented in the R package rMVP (v1.0.8) (Yin et al., 2021). FarmCPU was employed due to its robust capacity to enhance statistical power while strictly controlling for false positives. This is achieved through an iterative procedure that alternates between a Fixed Effect Model (FEM) and a Random Effect Model (REM), effectively decoupling the confounding effects of kinship from candidate markers and preventing the overfitting issues common in traditional stepwise regression.

Two types of resistance-related traits were analyzed: survival status (a binary trait: 0 for dead, 1 for survived) and survival time (a quantitative trait, h). To account for population stratification and individual morphological variation, the first five principal components (PC1-PC3), along with body length (BL) and body weight (BW), were incorporated as fixed-effect covariates and the GRM as a random effect. The FEM stage of the FarmCPU model is expressed as:

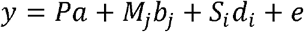

where y is the vector of phenotypic values; P represents the matrix of fixed effects (PC1–PC3, BL and BW); *a* is the vector of corresponding coefficients; *M*_*j*_ denotes the genotypes of the j-th pseudo-quantitative trait nucleotides (pseudo-QTNs) used as fixed effects; *b*_*j*_ is the effect size of the j-th pseudo-QTNs; *S*_*i*_ is the genotype of the i-th SNP marker being tested; *d*_*i*_ is its effect size; and *e* is the vector of residual errors. In the subsequent REM stage, the kinship matrix is iteratively updated based on the identified pseudo-QTNs to further refine the random polygenic effects.

To account for multiple testing, genome-wide significance thresholds were determined using the Bonferroni correction. The genome-wide significance threshold was set at p < 0.05/N, and the suggestive threshold at p < 1.0/N (where N is the total number of filtered SNPs). The GWAS results were visualized using Manhattan plots to illustrate the chromosomal distribution of −log10(p) values and quantile-quantile (Q-Q) plots to evaluate model fitting and identify potential inflation or deflation of p-values.

### 2.5. Genetic Differentiation Analysis (***F***_ST_)

To validate the genomic regions identified by GWAS and further characterize the genetic divergence between the two phenotypic groups, the fixation index (*F*_ST_) was calculated. The 300 individuals were divided into two populations: the resistant group (n=150) and the susceptible group (n=150). Genome-wide *F*_ST_ values were estimated using VCFtools (v0.1.16) (Danecek et al., 2011) based on the Weir and Cockerham (1984) estimator. A sliding-window approach was implemented to smooth the *F*_ST_ values across the genome, with a window size of 100 kb and a step size of 10 kb.

Windows containing fewer than 10 SNPs were excluded to minimize stochastic noise. The *F*_ST_ values were then Z-transformed (*Z-F*_ST_), and the genomic regions falling within the top 1% of the *F*_ST_ distribution were identified as highly differentiated regions (HDRs). Finally, the overlapping regions between the GWAS significant signals and the HDRs were prioritized as the most robust candidate genomic intervals associated with *Vibrio harveyi* resistance in *P. diacanthus*.

### 2.6. Candidate gene annotation and functional analysis

The locations of significant SNPs and InDels across various genomic regions— including intergenic, intronic, and exonic regions—were identified using the General Feature Format (GFF) file of the *P. diacanthus* reference genome (GCA_028641955.1). To identify candidate genes, we scanned the genomic neighborhoods of all markers that reached genome-wide significance for both survival status and survival time phenotypes. To ensure the comprehensive capture of proximal candidate genes and potential regulatory elements, a genomic search window of ±50 kb flanking the lead significant variants was established, a distance consistent with the typical linkage disequilibrium (LD) decay observed in teleost genomes. The identified sequences and gene models were functionally characterized through sequence homology searches using NCBI BLAST programs (Altschul et al., 1990) against the NCBI NR, NT, and UniProt databases.

### 2.7. Descriptive statistics and heritability estimation

Descriptive statistics for morphological traits (BL and BW) and survival parameters (survival time and status) were calculated using R (v4.3.1). To evaluate the genetic basis of *V. harveyi* resistance, the narrow-sense heritability (*h*^*2*^) for both traits was estimated using the Genomic Best Linear Unbiased Prediction (GBLUP) method implemented in the HIBLUP (v1.3.1) software (Yin et al., 2023). A Linear Mixed Model (LMM) was employed to partition the phenotypic variance into additive genetic and residual components. The model is expressed as follows:

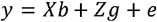

where *y* is the vector of phenotypic observations (survival time or survival status); *b* is the vector of fixed effects, including the top three principal components (PCs), and body weight (BW); *g* is the vector of random additive genetic effects, assumed to follow 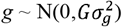, where *G* is the genomic relationship matrix (GRM) constructed from all high-quality SNPs and 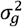 is the additive genetic variance; *X* and *Z* are the incidence matrices for fixed and random effects, respectively; and *e* is the vector of residual errors,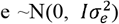. For the ST trait, heritability was estimated directly on the quantitative scale.

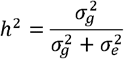

For the SS trait, heritability was first estimated on the observed scale and subsequently transformed to the liability scale using the following formula to account for the prevalence of mortality in the population:

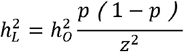

where 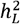 is the heritability on the liability scale, 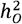is the heritability on the observed scale, *p* is the proportion of affected individuals (mortality rate) in the experimental population, and *z* is the height of the normal curve ordinate at the threshold corresponding to fraction *p*.

Phenotypic correlations among body weight (BW), survival time (ST), and survival status (SS) were evaluated using Pearson’s correlation coefficients via the “ggpubr” package in R. To estimate the genetic correlations *r*_*g*_ between these traits, a multi-trait linear mixed model was implemented using the sommer package (v4.3.0) in R. The genetic correlation was calculated as follows:

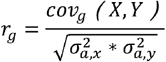

where *cov*_*g*_(*X, Y*) is the additive genetic covariance between traits X and Y, and 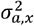 and 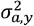 are the additive genetic variances for each trait, respectively. In the sommer framework, the first three principal components (PCs) were included as fixed-effect covariates. An unstructured covariance matrix was specified to simultaneously estimate the additive genetic variances and covariances based on the genomic relationship matrix (GRM). This multi-trait approach allowed for the partition of phenotypic correlations into genetic and environmental components, providing insights into the shared genetic basis between growth and *V. harveyi* resistance in *P. diacanthus*.

## 3. Results

### 3.1. Mortality after *V. harveyi* challenge

Throughout the 21-day monitoring window, all 45 individuals in the sham-injected control cohort remained symptom-free and achieved a 100% survival rate, verifying that the physical inoculation procedure did not induce any confounding non-specific mortality. Conversely, the *P. diacanthus* specimens within the infected treatment group displayed distinct clinical manifestations characteristic of *V. harveyi* pathogenesis. The diseased fish displayed multi-organ pathologies, including severe fin and skin erosion, exophthalmia (eye hemorrhage), and extensive internal hemorrhaging of the visceral organs (Figure 1). The mortality kinetics followed an acute infection pattern (Figure 2). Mortality was first recorded at 15 hours post-infection (hpi) and increased significantly starting at 19 hpi. The peak mortality period occurred between 19 and 32 hpi, with the median lethal time (LT_50_) reached at 32 hpi. After this peak, the mortality rate gradually plateaued. Upon the conclusion of the infection trial, a total of 240 individuals succumbed to the pathogen and were designated as the susceptible cohort, whereas 210 specimens successfully survived in a healthy state and were categorized as the resistant group. The cumulative mortality rate was 53.33%, which provided an ideal phenotypic distribution for subsequent genetic association studies.

**FIGURE 1.**
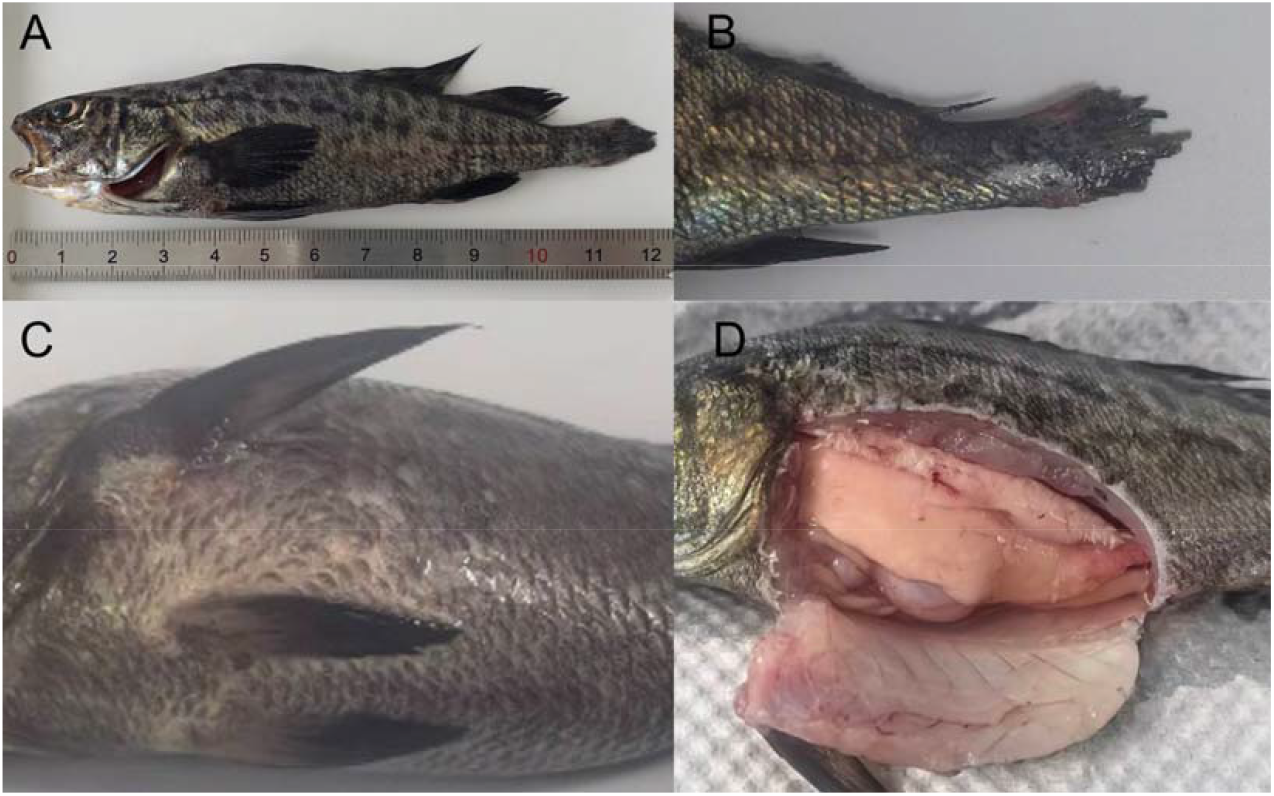
Clinical symptoms and pathomorphological changes in *P. diacanthus* challenged with *Vibrio harveyi*. (A) Whole-body view of a morbid fish displaying darkened body coloration and mouth agape; (B) severe caudal fin erosion with frayed rays and localized hyperemia; (C) lateral-ventral skin congestion and minor hemorrhage; (D) visceral view of dissected diseased fish showing multi-organ congestion and prominent liver blanching (whitened liver).

**FIGURE 2.**
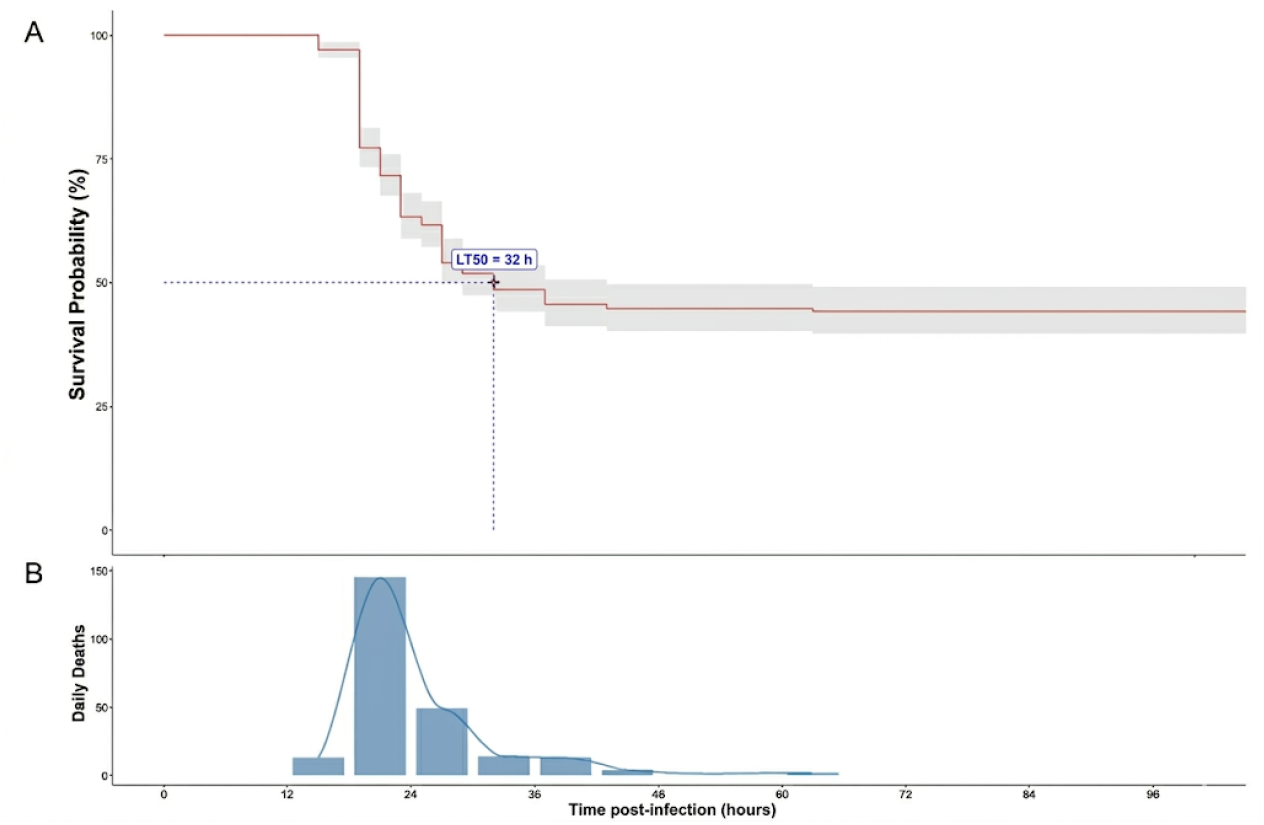
Mortality kinetics of *P. diacanthus* juveniles following *V. harveyi* challenge. (A) Kaplan-Meier survival curve showing the survival probability over 96 hours post-infection (hpi). The red line represents the survival rate, while the light gray shaded area indicates the 95% confidence interval. The blue dashed lines highlight the median lethal time (LT_50_), which occurred at 32 hpi. (B) Distribution of mortality across the challenge period. The blue bars represent the number of deaths at different time intervals, and the solid curve represents the kernel density estimation of the mortality rate, indicating a clear peak between 19 and 32 hpi.

### 3.2. Sequencing data and distribution of SNPs

Whole-genome resequencing of 300 individuals (150 resistant and 150 susceptible) generated approximately 3.3 Tb of raw data. After quality control, the average clean data per individual reached 11 Gb, with a mean sequencing depth of 18.4x. The high quality of the sequencing data provided sufficient resolution for accurate variant calling and downstream genetic analysis.

By aligning the clean reads to the *P. diacanthus* male reference genome, a total of 3,743,916 SNPs and 701,829 InDels were initially identified. Following stringent filtering, a high-quality subset of 2,041,727 SNPs and 601,790 InDels was retained. These markers were comprehensively distributed across all 24 chromosomes (Figure 3). As illustrated in the marker density map (Figure 3), SNP density was calculated within 1Mb window sizes (Figure 3A), while InDel density was assessed within 0.1Mb window sizes (Figure 3B). The distribution profiles revealed that the variants covered the entire genome with high density, although variations in density were observed across different chromosomal regions. Notably, certain regions on chromosomes such as Chr12 and Chr23 exhibited higher marker concentrations. This high-density marker set ensured robust genomic coverage for subsequent association mapping and selection sweep analyses.

**FIGURE 3.**
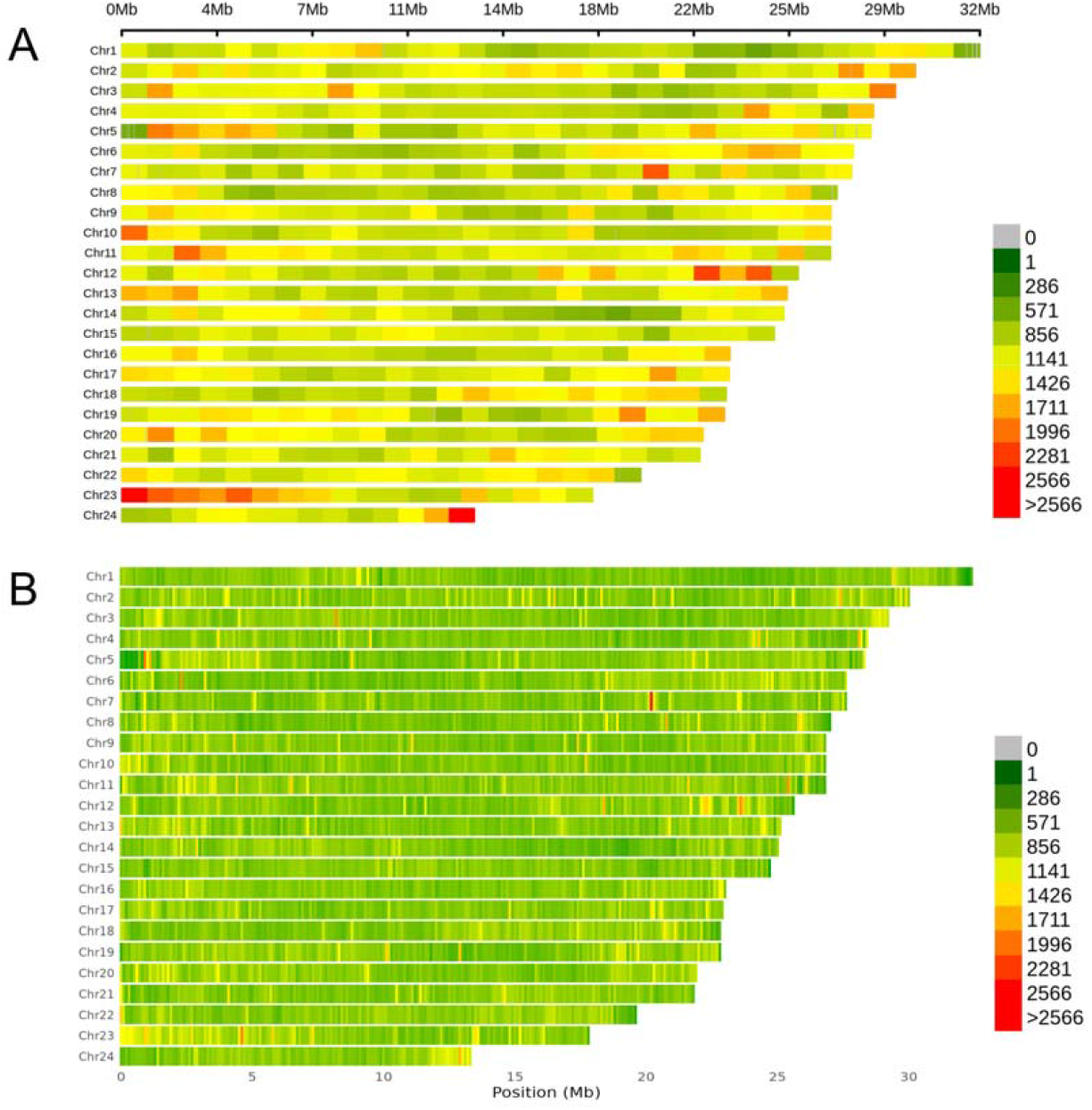
Genomic distribution and density of SNPs and InDels across the 24 chromosomes of *P. diacanthus*. (A) SNP density calculated within 1 Mb window sizes. (B) InDel density calculated within 0.1 Mb window sizes. The horizontal axis represents the chromosomal position (Mb), and the vertical axis represents the chromosome number (Chr1–Chr24). The color scale from green to red indicates increasing marker density (number of variants per window).

### 3.3. Analysis of population structure and genetic relatedness

To investigate the population structure of the experimental *P. diacanthus*, principal component analysis (PCA) and Neighbor-Joining (NJ) phylogenetic tree analysis were performed. The PCA results, based on both SNPs (Figure 4A) and InDels (Figure 4B), revealed that the population primarily consisted of a single major cluster. While some individuals exhibited slight dispersion at the periphery of the main cluster, the resistant (alive) and susceptible (dead) individuals were uniformly distributed throughout the PCA space without distinct phenotypic clustering. A substantial proportion of the total genetic variation was captured by the leading axes, with PC1 and PC2 accounting for 45.2% and 32.22% of the variance, respectively. Consistently, the genome-wide NJ phylogenetic tree (Figure 4C) showed that resistant and susceptible individuals were highly intermingled across different branches, further confirming the absence of significant population stratification between the two phenotypic groups. To account for subtle sub-structures in the subsequent GWAS, the top principal components were incorporated as fixed effects, and the genomic relationship matrix (GRM) was included as a random effect. As illustrated in the kinship heatmap (Figure 4D), low degree of cryptic relatedness was evident within the study cohort, as the vast majority of pairwise kinship coefficients fell well beneath the 0.2 threshold. These findings demonstrate that the population structure was well-controlled, providing a reliable basis for identifying true genetic associations with *V. harveyi* resistance.

**FIGURE 4.**
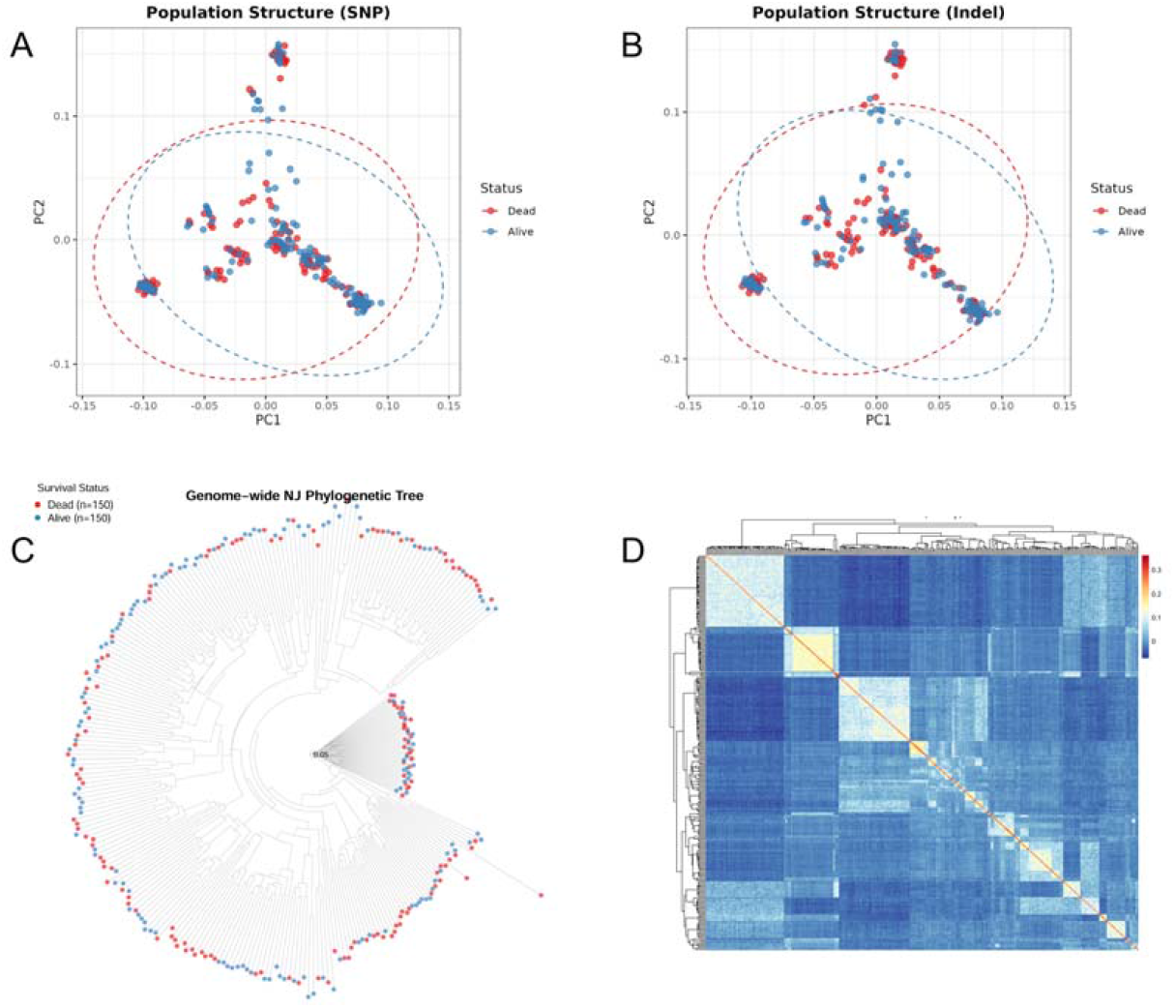
Population structure and genetic relatedness analysis of the 300 *P. diacanthus* individuals. (A-B) Principal component analysis (PCA) plots based on SNPs and InDels, respectively. Red dots represent dead (susceptible) individuals, and blue dots represent alive (resistant) individuals. (C) Genome-wide Neighbor-Joining (NJ) phylogenetic tree showing the genetic relationships among all individuals. (D) Heatmap of the genomic relationship matrix (GRM) depicting pairwise kinship coefficients. The color scale from blue to red represents increasing genetic similarity.

### 3.4. Genome-wide association study

To identify genomic regions associated with *V. harveyi* resistance, GWAS was performed for both survival hours (ST) and survival status (SS) using the FarmCPU model. The quantile-quantile (Q-Q) plots (Figure 5A–D) show that the observed *−log*_*10*_ (P) values followed the expected null distribution for most markers but deviated significantly at the tail, indicating that the model effectively controlled for population structure and successfully identified true association signals. Significant association signals were consistently detected across both phenotypes and both marker types (SNPs and InDels). A major genomic cluster exceeding the genome-wide Bonferroni significance threshold (p < 0.05/N) was identified on Chromosome 4 (Chr4) for both ST and SS (Figure 5A, C). Additional significant or suggestive signals were observed on Chr5, Chr12, Chr15, and Chr16. The lead SNP on Chr4 (Chr4_12706015 bp) exhibited the strongest association, suggesting the presence of a primary quantitative trait locus (QTL) in this region. To further validate the predictive power of the identified markers, genotype-phenotype heatmaps were constructed for the top 50 SNPs (Figure 5E) and top 50 InDels (Figure 5F). The heatmaps revealed a clear segregation between the resistant (survived) and susceptible (dead) groups based on their multi-locus genotypes. Specifically, individuals that survived the challenge shared distinct homozygous or heterozygous genotypes at these top-ranking loci, whereas susceptible individuals predominantly carried alternative alleles. The high consistency between the GWAS signals for survival hours and status, coupled with the clear clustering in the heatmap analysis, underscores the robustness of the identified genomic regions in conferring resistance to *V. harveyi*.

**FIGURE 5.**
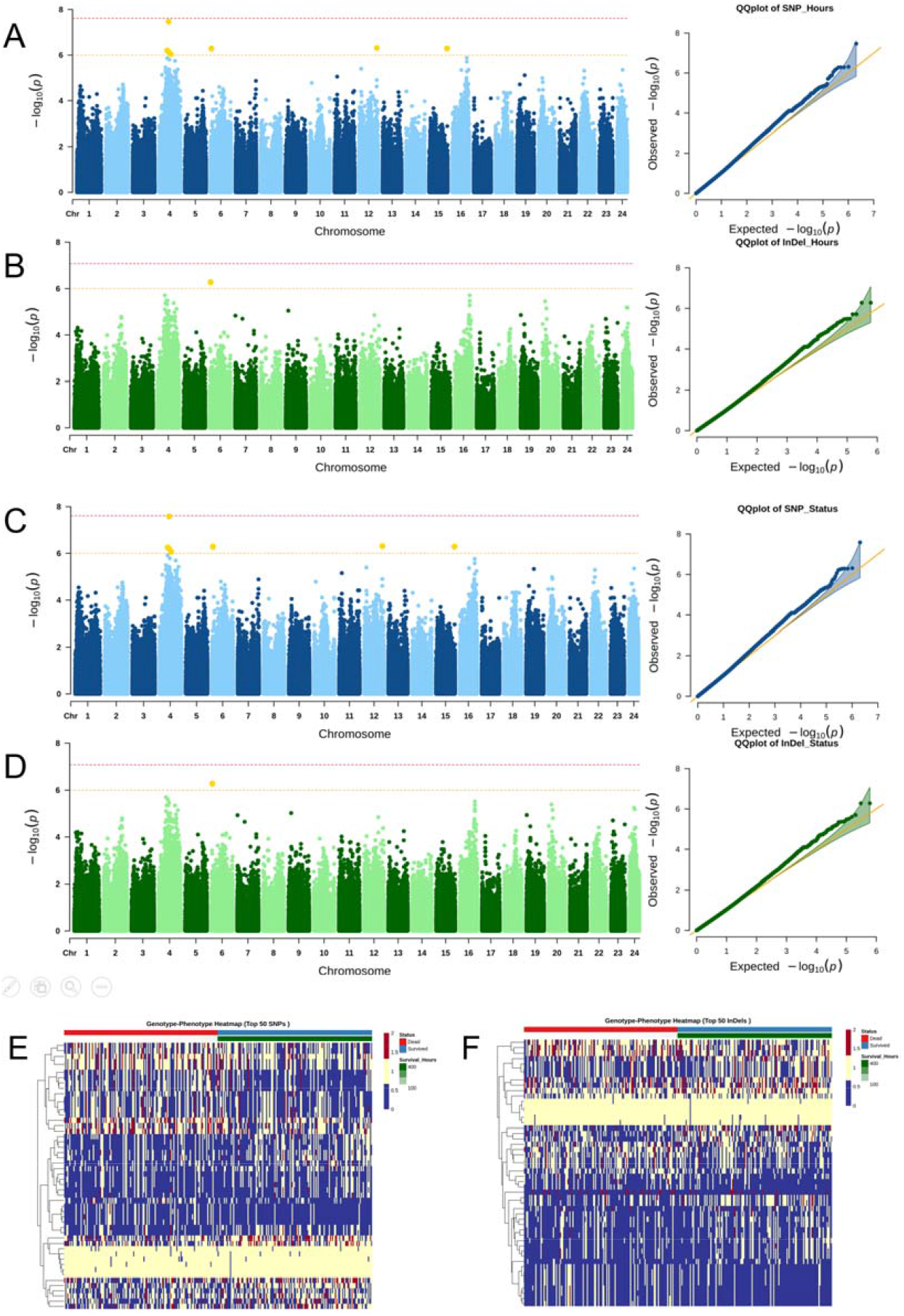
Genome-wide association study (GWAS) of *V. harveyi* resistance in *P. diacanthus*. (A-D) Manhattan plots (left) and Q-Q plots (right) for SNP-based survival hours (A), InDel-based survival hours (B), SNP-based survival status (C), and InDel-based survival status (D). The horizontal red and orange dashed lines represent the genome-wide significance (p < 0.05/N) and suggestive (p < 1.0/N) thresholds, respectively. (E-F) Genotype-phenotype heatmaps for the top 50 SNPs (E) and top 50 InDels (F). The top color bar represents the phenotypic status (red: dead; green: survived), and the rows represent individual markers.

### 3.5. Genetic Differentiation Analysis (*F*_ST_)

To further validate the candidate genomic regions and identify loci under divergent selection between the two phenotypic groups, we calculated the genome-wide fixation index (*F*_ST_) between the resistant (n = 150) and susceptible (n = 150) populations. The genome-wide *F*_ST_ distribution (Figure 6) revealed several regions with significant genetic differentiation, suggesting that these loci may be associated with the observed survival variance. Importantly, several *F*_ST_ peaks exhibited strong spatial co-localization with the significant GWAS signals identified previously. Notably, a prominent cluster of high *F*_ST_ values was observed on Chromosome 4 (Chr4), coinciding with the major QTL region detected in the Manhattan plots for survival status and hours. Additional differentiation signals were detected on Chr6, Chr12, and Chr16, which also showed suggestive or significant associations in the GWAS analysis. The convergence of results from both association mapping and genetic differentiation analysis provides robust evidence for the role of these genomic intervals, particularly the regions on Chr4, in conferring resistance to *V. harveyi* in *P. diacanthus*.

**FIGURE 6.**
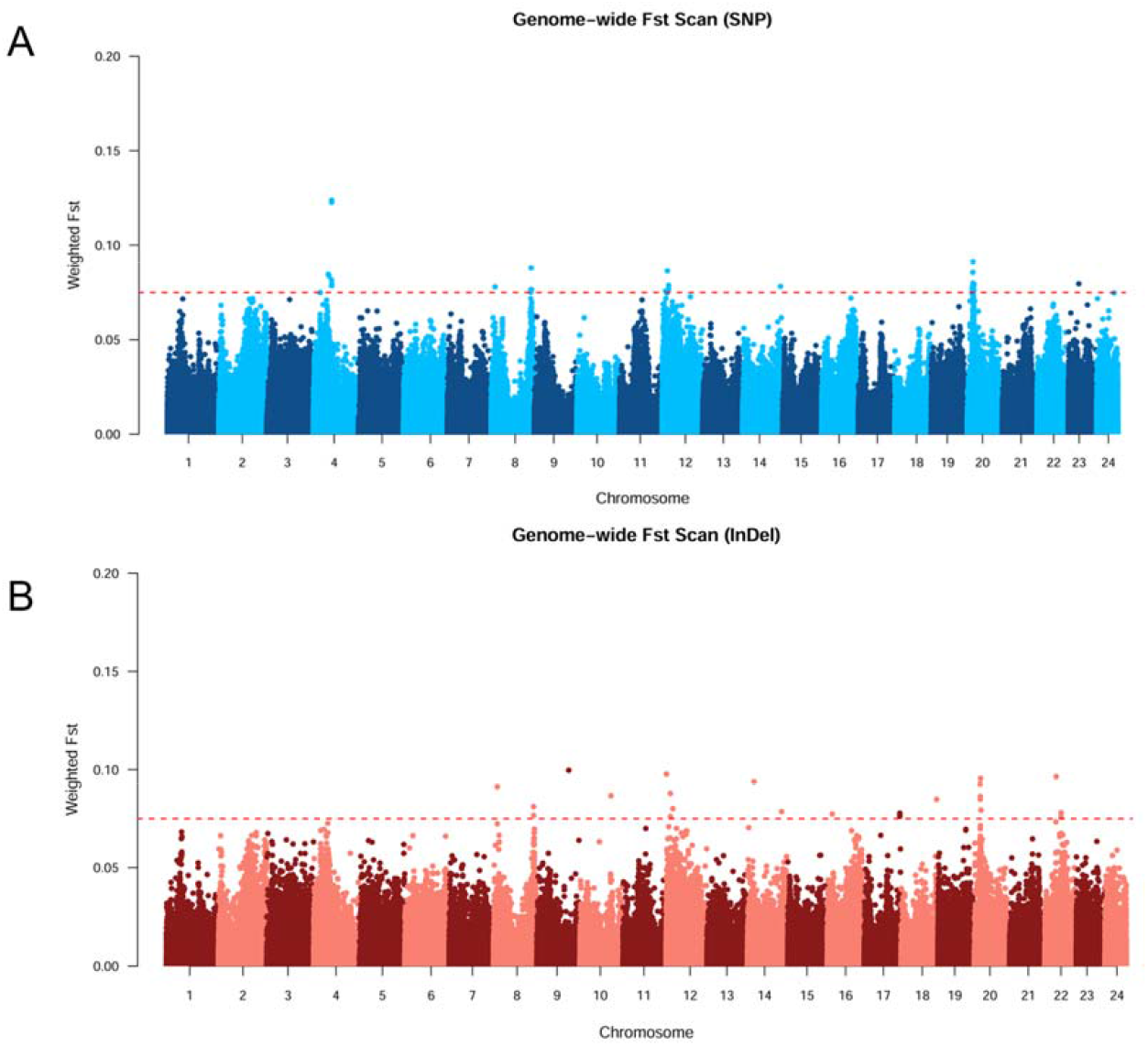
Genome-wide genetic differentiation (*F*_ST_) between resistant and susceptible *P. diacanthus* groups. (A) Manhattan plot of *F*_ST_ values calculated based on SNP markers. (B) Manhattan plot of *F*_ST_ values calculated based on InDel markers. The horizontal axis represents the 24 chromosomes, and the vertical axis indicates the weighted *F*_ST_ values. The red dashed line denotes the top 1% (or 0.5%) significance threshold. Peaks on chromosome 4 show strong spatial co-localization with GWAS signals.

### 3.6. Descriptive statistics and heritability estimation

A comprehensive overview of the statistical parameters describing growth performance within this experimental cohort is presented in Table 1. This phenotypic trend was further visualized in the distribution plots (Figure 7), providing a comprehensive overview of the population’s morphological characteristics during the *V. harveyi* challenge. Genomic heritability ( *h*^*2*^) and its associated variance components were estimated using the GBLUP method (Table 2). The narrow-sense heritability for survival status (SS) was 0.237±0.185, while survival time (ST) exhibited a higher heritability of 0.374±0.29. These results indicate that resistance to *V. harveyi* in *P. diacanthus* is under moderate genetic control, thereby underscoring a substantial scope for achieving rapid genetic gains via systematic selection frameworks. The genetic (*r*_*g*_) and phenotypic (*r*_*p*_) correlations among growth and resistance traits are presented in Table 3. A near-perfect correlation (*r*_*g*_ = 0.999, *r*_*p*_ = 0.999) was observed between SS and ST, demonstrating that these two metrics of disease resistance are genetically and phenotypically synonymous.

**TABLE 1.** Descriptive statistics of growth traits.

| Mean | Total | Non-survivors | survivors |
| --- | --- | --- | --- |
| BL (cm) | $12.79 \pm 1.39$ | $12.02 \pm 1.01$ | $13.55 \pm 1.39$ |
| BD (g) | $38.38 \pm 9.54$ | $36.80 \pm 8.23$ | $39.95 \pm 10.49$ |

**TABLE 2.**
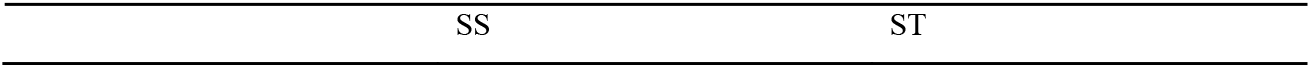
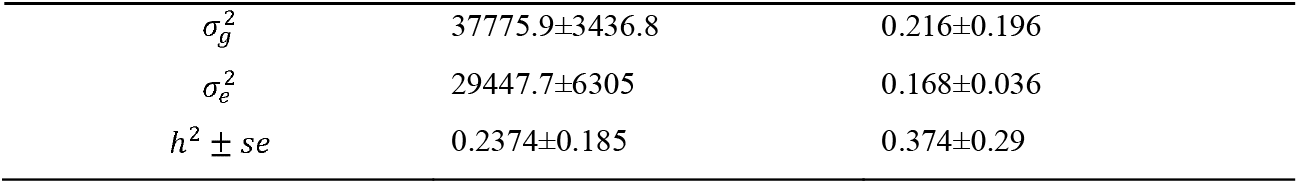
Variance components and genomic heritability (*h*^*2*^) estimates.

| SS | ST |
| --- | --- |

|  |  |  |
| --- | --- | --- |
| $\sigma_g^2$ | 37775.9±3436.8 | 0.216±0.196 |
| $\sigma_e^2$ | 29447.7±6305 | 0.168±0.036 |
| $h^2 \pm se$ | 0.2374±0.185 | 0.374±0.29 |

**TABLE 3.** Genetic (*r*_*g*_) and phenotypic (*r*_*p*_) correlations among growth and resistance traits.

|  | SS | ST | BW |
| --- | --- | --- | --- |
| SS | 1 | 0.999 | 0.0289 |
| ST | 0.999 | 1 | 0.0183 |
| BW | 0.1714 | 0.1662 | 1 |

**FIGURE 7.**
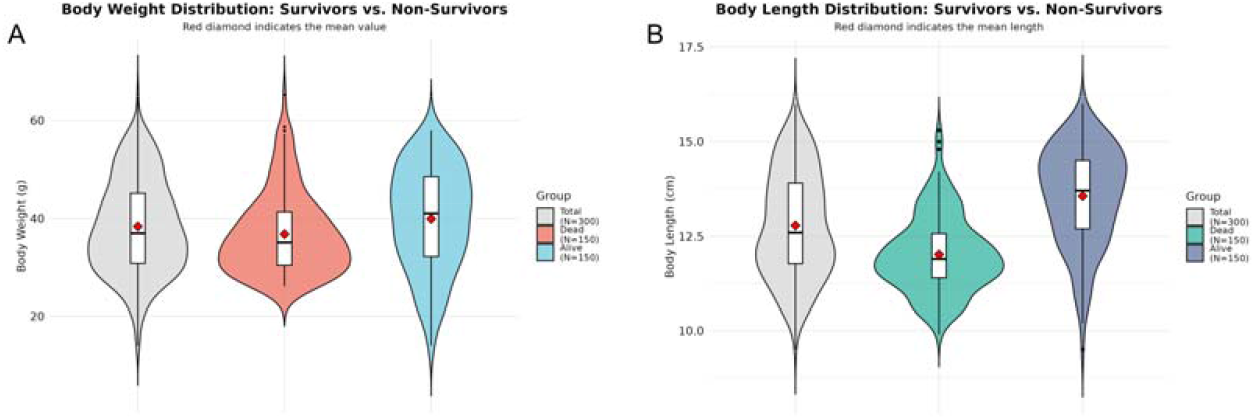
Comparison of body length (BL) (A) and body weight (BW) (B) distribution between the surviving and dead groups

In contrast, the relationship between growth and resistance was characterized by very low correlations. The genetic correlations between BW and resistance traits were 0.0289 (with SS) and 0.0183 (with ST). The phenotypic correlations were slightly higher but remained low (0.1714 with SS and 0.1662 with ST). These findings confirm that growth rate and *V. harveyi* resistance are genetically independent in *P. diacanthus*, implying that breeding for faster growth will not negatively impact the innate disease resistance of the population.

### 3.7. Candidate gene annotation and functional analysis

To screen for putative functional determinants governing defense against *V. harveyi* in a methodical manner, genomic regions ±50 kb flanking the top 7 significant SNPs and InDels were annotated (Table 4). This analysis identified a group of high-confidence candidates involved in RNA stability regulation, transcriptional control, and endocytosis. On Chromosome 4 (Chr4), the most prominent association signal (Top1) was localized within the exon of *elavl4* (ELAV-like family member 4), suggesting that a functional polymorphism in this RNA-binding protein may influence host resistance by modulating mRNA stability or translation. This chromosome also harbored several critical gene clusters. Cluster A contained the gene *rorb* (Retinoic acid receptor-related orphan receptor beta), which was associated with an exon variant. Cluster B included *cbpc6* (CBP/p300-interacting transactivator 6) situated near an intron variant. These findings indicate that Chr 4 represents a major genomic reservoir for resistance, likely acting through the coordination of gene transcription and post-transcriptional regulation. Furthermore, on Chromosome 15 (Chr15), a significant locus was identified in the intergenic region adjacent to *epn4* (Epsin-4), a protein primarily involved in clathrin-mediated endocytosis and membrane trafficking. On Chromosome 12 (Chr12), a strong association signal was found adjacent to *Nibea_GLEAN_10013770*; although this gene currently lacks a Swissprot description, its proximity to a highly significant marker suggests it may represent a novel or species-specific candidate for disease resistance in *P. diacanthus*.

**TABLE 4.** Summary of top candidate genes associated with *V. harveyi* resistance.

| Traits | ID | Type | Chr | Position (bp) | P(value) | beta | Pve(%) | MAF | Allele | Association | Variant | Gene |
| --- | --- | --- | --- | --- | --- | --- | --- | --- | --- | --- | --- | --- |
| SS/ST | 1 | SNP | 4 | 12706015 | 3.41E-08 | -87.9565 | 3.242247 | 0.193 | T/C | Significant | Exon | *elavl4* |
| SS/ST | 2 | SNP | 12 | 23780687 | 4.89E-07 | 681.1854 | 4.240472 | 0.498 | G/A | Suggestive | Intergenic |  |
| SS/ST | 3 | SNP | 15 | 23978134 | 5.14E-07 | 680.1158 | 4.224641 | 0.498 | G/A | Suggestive | Intergenic | *epn4* |
| SS/ST | 4 | Indel | 6 | 1199929 | 5.25E-07 | 679.5684 | 4.217819 | 0.498 | A/ATTGC | Suggestive | Intergenic |  |
| SS/ST | 5 | SNP | 4 | 10842456 | 6.28E-07 | -65.9076 | 3.855096 | 0.362 | A/G | Suggestive | Exon | *rorb* |
| SS/ST | 6 | SNP | 4 | 12585128 | 7.19E-07 | -59.6439 | 4.045466 | 0.433 | T/C | Suggestive | Intron | *cbpc6* |
| SS/ST | 7 | SNP | 4 | 15581089 | 9.06E-07 | 96.07473 | 1.78485 | 0.123 | A/G | Suggestive | Intergenic |  |

GO and KEGG enrichment analyses were performed to elucidate the functional landscape of these candidate genes. GO analysis revealed significant enrichment in terms related to the “cell surface receptor signaling pathway”, “activation of immune response”, and “defense response to bacterium” (Figure S1A). Consistently, KEGG pathway analysis showed that these candidate genes were significantly represented in the Toll-like receptor (TLR) signaling pathway, the MAPK signaling pathway, and Apoptosis (Figure S1B). The integration of specific candidates such as *elavl4* and *rorb* with the identified immune-related pathways suggests that resistance to *V. harveyi* in *P. diacanthus* involves a complex regulatory network encompassing mRNA metabolic processes and classical innate immune signaling.

## 4. Discussion

In the present study, we conducted the first comprehensive genome-wideassociation study (GWAS) integrated with genetic differentiation analysis (*F*_*st*_) to systematically unravel the genetic architecture of *V. harveyi* resistance in the blackspotted croaker (*Protonibea diacanthus*). Our findings demonstrate that vibriosis resistance in this species is a polygenic trait under moderate genomic control. Notably, we identified a pivotal quantitative trait locus (QTL) on chromosome 4 and prioritized a suite of high-confidence candidate genes, including *nop56, nop58*, and *utp5*. These results address a fundamental knowledge gap in *P. diacanthus* genetics and establish a robust molecular framework for future marker-assisted selection (MAS) and genomic selection (GS) programs aimed at enhancing disease resistance in the sciaenid industry.

### 4.1. Heritability of Disease Resistance and Its Implications for Breeding

In the present study, genomic heritability (*h*^2^) for *V. harveyi* resistance in *P. diacanthus* was estimated utilizing a GBLUP framework. The narrow-sense heritability was 0.374 ± 0.29 for survival time (ST) and 0.237 ± 0.185 for survival status (SS) (Table 2). These values classify resistance to vibriosis as a trait under moderate genetic control, providing robust evidence that selective breeding programs can effectively enhance the innate immunity of this species.

The heritability estimates obtained here align closely with those reported in other sciaenids and economically important marine teleosts. For instance, heritability for *V. harveyi* resistance in *N. albiflora* was reported between 0.28 and 0.35 (Luo et al., 2021). Similar ranges have been observed in *Cynoglossus semilaevis*, where *h*^2^ was estimated at 0.28–0.45 (Li et al., 2019; Hu et al., 2020). Furthermore, our results are consistent with estimates for the *Plectropomus leopardus* (*h*^2^ = 0.32-0.41; Lu et al., 2023) and the *Hippocampus erectus* (*h*^2^≈0.28; Li et al., 2024). In the large yellow croaker (*Larimichthys crocea*), comparable moderate heritability has been documented for resistance against both *Cryptocaryon irritans* (0.32-0.46; Zhao et al., 2021) and *Pseudomonas plecoglossicida* (0.31-0.40; Wan et al., 2019). The consistency of these findings across diverse taxa underscores the general feasibility of genetic improvement for bacterial disease resistance in aquaculture.

A notable observation in this study was that the heritability for survival time (ST) surpassed that of survival status (SS), a trend mirroring recent findings in leopard coral grouper (Lu et al., 2023) and Pacific oyster (*Crassostrea gigas*) (Yang et al., 2024). This disparity is likely attributed to the nature of the phenotypic data; as a continuous quantitative trait, ST captures nuanced variation regarding the progression of mortality, whereas the binary nature of SS (survived/dead) provides a more simplified phenotypic snapshot. Consequently, ST may offer superior statistical power for resolving the genetic architecture of disease resistance. Based on this, we propose that survival time should be prioritized as a primary phenotypic indicator in future genomic evaluation and selection programs for *P. diacanthus*.

Perhaps the most significant finding for industrial application is the near-zero genetic correlation between growth performance and disease resistance. Our analysis revealed that genetic correlations between body weight (BW) and resistance traits (SS and ST) were negligible (0.0289 and 0.0183, respectively; Table 3). This genetic independence has also been reported in yellow drum (Luo et al., 2021) and Pacific oyster (Yang et al., 2024). From a breeding perspective, this is an ideal scenario: it demonstrates that rapid growth and enhanced vibriosis resistance are not mutually exclusive. Therefore, breeders can simultaneously implement multi-trait selection strategies to develop elite *P. diacanthus* lines that combine superior growth rates with high survival under *V. harveyi* challenge, without the risk of negative genetic antagonism.

### 4.2. Cross-validation of GWAS and F_ST_ Analysis Enhances Candidate Region Reliability

In the present study, GWAS unveiled a highly significant association signal on chromosome 4 (Chr4_12706015, p = 3.41 × 10^-8^), which was consistently associated with both survival time (ST) and survival status (SS) (Table 4). Concurrently, genome-wide *F*_ST_ analysis between the resistant and susceptible groups manifested highly differentiated regions (HDRs), with the most prominent peak also localized on chromosome 4 (Figure 6). This spatial co-localization of GWAS and *F*_ST_ signals provides robust cross-validation for the candidate region, substantially enhancing its biological reliability.

*F*_ST_ is a classical measure of allele frequency divergence between populations, reflecting the genomic impact of natural or artificial selection. When GWAS signals overlap with *F*_ST_ peaks, it indicates that the genomic region is not merely statistically associated with the phenotype but has also undergone substantial allele frequency shifts between the phenotypic extremes (resistant vs. susceptible). Such a synergistic methodology has become a benchmark in recent aquaculture genomics for pinpointing high-confidence loci. In the yellow drum, GWAS-significant SNPs were found to be localized within regions of elevated *F*_ST_ (Luo et al., 2021). Similarly, in the leopard coral grouper, cross-validation strategies were successfully utilized to confirm candidate regions for *V. harveyi* resistance (Lu et al., 2023). Furthermore, in the Pacific oyster, the combination of GWAS and *F*_ST_ analysis effectively narrowed candidate intervals for vibriosis resistance (Yang et al., 2024). By employing this multi-method validation, we significantly mitigate the risk of false-positive associations and bolster the credibility of the identified resistance-related loci in *P. diacanthus*.

### 4.3. Functional Annotation of Candidate Genes: From RNA Metabolism to Immune Regulation

The lead candidate genes identified in this study are primarily located within ±50 kb of the significant markers on chromosome 4 and chromosome 15, including *elavl4* (ELAV-like family member 4), *rorb* (Retinoic acid receptor-related orphan receptor beta), *cbpc6* (CBP/p300-interacting transactivator 6), and *epn4* (Epsin-4) (Table 4). Unlike traditional structural immune genes, these candidates represent a sophisticated regulatory network encompassing post-transcriptional processing, transcriptional modulation, and membrane trafficking.

On chromosome 4, *elavl4* (also known as HuD) stands out as a premier candidate due to its role as an RNA-binding protein. Proteins belonging to the ELAVL family are widely recognized for targeting AU-rich elements (AREs) situated inside the 3′ untranslated regions (UTRs) of downstream transcripts, thereby significantly increasing their mRNA stability and translational efficiency (Silvestri et al., 2022). In the context of *V. harveyi* infection, the rapid stabilization of mRNAs encoding cytokines or antimicrobial peptides is essential for an effective innate immune response. The identification of a significant exon variant in *elavl4* suggests that polymorphisms in this gene may dictate the speed and magnitude of the host’s “inflammatory surge” by modulating the half-life of key immune transcripts.

The presence of *rorb* and *cbpc6* on chromosome 4 further highlights the importance of transcriptional reprogramming in vibriosis resistance. RORB is a nuclear receptor that acts as a ligand-dependent transcription factor (Jetten et al., 2006). These factors are critical for orchestrating the cellular response to bacterial stress and maintaining immune homeostasis. Concurrently, on chromosome 15, *epn4* (Epsin-4) is involved in clathrin-mediated endocytosis, a process vital for the internalisation of cell-surface receptors and the uptake of bacterial toxins or pathogens (Sen et al., 2012).

Our functional enrichment analyses provide compelling evidence for this regulatory landscape. GO enrichment analysis (Figure. S1A) showed that the candidate genes are predominantly involved in “rRNA processing”, “mRNA processing”, and “Ribosome biogenesis”. These biological processes are fundamental for maintaining cellular homeostasis under stress. Specifically, the identification of *elavl4* (ELAV-like family member 4) as a top candidate aligns perfectly with the enriched GO terms such as “RNA binding” and “mRNA metabolic process”. ELAVL4 is a well-known RNA-binding protein that stabilizes target mRNAs, thereby ensuring the efficient translation of essential proteins during the host’s response to bacterial infection. Consistently, KEGG pathway analysis (Figure. S1B) revealed that the most significantly enriched pathways are “Ribosome biogenesis in eukaryotes”, “RNA transport”, “mRNA surveillance pathway”, and “Nucleocytoplasmic transport”. The prominence of the mRNA surveillance pathway is particularly significant; this system acts as a quality control mechanism to detect and degrade aberrant mRNAs, which is vital for the host to maintain a functional proteome during the rapid physiological changes induced by Vibriosis. The identification of transcriptional modulators like *rorb* and *cbpc6* further supports the importance of coordinated gene expression and RNA transport in the resistance phenotype. Although *epn4* (Epsin-4) is primarily associated with endocytosis, its role in membrane trafficking may complement the efficient transport of signaling molecules within the enriched “Nucleocytoplasmic transport” framework (Akira et al., 2004; Kawai & Akira, 2010; Zhang et al., 2022).

In summary, the convergence of RNA-binding proteins (*elavl4*), transcriptional modulators (*rorb, cbpc6*), and endocytic factors (*epn4*) suggests that resistance to *V. harveyi* in *P. diacanthus* is not governed by a single “immune switch” but rather by a multi-layered regulatory architecture. The convergence of genomic association signals within pathways related to RNA metabolism and ribosome assembly represents a novel insight into the genetic basis of disease resistance in *P. diacanthus*. These findings suggest that resistant individuals may possess a more efficient or robust system for RNA quality control and protein synthesis, allowing them to rapidly mobilize cellular resources and maintain physiological stability when challenged by *V. harveyi*.

### 4.4. Comparison with Related Species: Species-Specific Genetic Architecture of Disease Resistance

The major QTL identified in this study is localized on chromosome 4, whereas in the closely related *N. albiflora*, the primary QTL for *V. harveyi* resistance was mapped to chromosome 14 (Luo et al., 2021). Subsequent integrative GWAS and eQTL analyses in the yellow drum identified candidate genes directly involved in canonical immune signaling, such as SOCS1, CLEC16A, CIITA, and PRKCB (Huang et al., 2024). In contrast, our top candidates in *P. diacanthus*—namely *elavl4, rorb*, and *cbpc6*—are primarily associated with RNA stability, transcriptional modulation, and mRNA metabolic processes. This discrepancy strongly suggests that, despite their phylogenetic proximity and shared pathogen pressure, the genetic architecture of *V. harveyi* resistance has diverged significantly between these two sciaenid species.

Such species-specific genetic architectures are a recurrent theme in fish disease resistance research. In the large yellow croaker (*Larimichthys crocea*), GWAS for resistance to *Cryptocaryon irritans* identified genes involved in immune recognition and cell adhesion (Zhao et al., 2021), while resistance to *Pseudomonas plecoglossicida* was associated with pathways related to apoptosis and inflammatory responses (Wan et al., 2019). Similarly, in rainbow trout (*Oncorhynchus mykiss*), resistance to *Flavobacterium columnare* is governed by QTL across multiple chromosomes involving mucosal immunity and immune signaling (Fraslin et al., 2022). In the common carp (*Cyprinus carpio*), resistance to koi herpesvirus (KHV) involves diverse mechanisms, including antiviral immunity and programmed cell death (Jia et al., 2020; Palaiokostas et al., 2018). Specific genomic loci governing antiviral defense cascades have been implicated in conferring tolerance to white spot syndrome virus (WSSV) within invertebrate models, such as the Pacific white shrimp (*Litopenaeus vannamei*) (Medrano-Mendoza et al., 2023). Collectively, these findings underscore that different aquatic species have evolved distinct genetic strategies to cope with similar microbial challenges.

One plausible explanation for the unique findings in *P. diacanthus* is that this species may rely more heavily on the fine-tuning of post-transcriptional regulation to mount an effective defense. By modulating the stability of immune-related mRNAs (via *elavl4*) and orchestrating rapid transcriptional reprogramming (via *rorb* and *cbpc6*), *P. diacanthus* may achieve a more flexible or robust response compared to species that depend predominantly on fixed immune signaling cascades. However, these strategies are not necessarily mutually exclusive. In the yellow drum, eQTL analyses also identified genes involved in RNA processing and translational regulation (Huang et al., 2024), suggesting that the post-transcriptional control of immune responses might be a conserved but differentially emphasized mechanism across *sciaenids*. Future comparative functional studies are warranted to further elucidate the evolutionary trade-offs and functional nuances of these candidate genes across diverse fish taxa.

### 4.5 Comparison with GWAS Studies in Other Aquatic Species

Extending the comparison to a broader spectrum of aquatic species further contextualizes the unique genomic landscape of *P. diacanthus*. In the Pacific oyster (*C. gigas*), GWAS for resistance to *V. alginolyticus* and general vibriosis identified candidate genes predominantly involved in immune recognition (e.g., C-type lectins) and apoptosis regulation (Yang et al., 2022; Yang et al., 2024). Similarly, in Atlantic salmon (*Salmo salar*), GWAS for sea lice (*Caligus rogercresseyi*) resistance highlighted QTL associated with immune signaling and skin barrier function (Correa et al., 2017; Robledo et al., 2019). While these studies emphasize classical host-pathogen interaction components, our findings in *P. diacanthus* point toward a more complex regulatory layer involving post-transcriptional and metabolic fine-tuning.

Notably, the top candidate genes identified in our study (*elavl4, rorb*, and *epn4*) are not classical immune effectors but rather essential regulators of RNA stability and intracellular trafficking. This highlights the possibility that in *P. diacanthus*, the efficiency of mRNA metabolic processes and surveillance may be a decisive factor in determining disease outcomes. Similar observations regarding the importance of cellular maintenance pathways have emerged in other taxa. In zebrafish (*Danio rerio*), high-throughput screenings revealed that genetic variation in RNA-processing and ribosome-related genes significantly contributes to differential susceptibility to environmental stressors, suggesting these pathways are fundamental determinants of individual resilience (Balik-Meisner et al., 2018; Wallis et al., 2022). Furthermore, in the yellowfin seabream (*Acanthopagrus latus*), integrated GWAS and transcriptomic analyses for *Streptococcus iniae* resistance identified candidates involved in protein synthesis and ribosome function (Pan et al., 2025).

These collective findings, along with our discovery of RNA-binding proteins and transcriptional modulators, suggest that cellular biosynthetic capacity and RNA quality control may be universal, yet underappreciated, factors influencing host resistance across diverse aquatic taxa. In *P. diacanthus*, the shift from traditional immune “switches” to a more nuanced RNA-centric regulatory network may represent an evolutionary adaptation for rapid physiological reconfiguration during acute bacterial challenge.

### 4.6. Limitations and Future Perspectives

While the present study provides novel insights into the genetic architecture of *V. harveyi* resistance in *P. diacanthus*, several limitations warrant acknowledgement to contextualize our findings. First, regarding sample size and statistical power, we employed an extreme phenotype sampling strategy, resequencing 300 individuals from a challenged population of 450. Although this approach effectively maximizes statistical power under budgetary constraints, a sample size of 300 remains relatively moderate for GWAS. This may limit the sensitivity required to detect minor-effect loci that contribute to the polygenic nature of disease resistance. Future investigations involving significantly larger cohorts and independent populations will be essential to capture the full spectrum of the genetic landscape. Second, the functional validation of prioritized candidate genes remains a critical next step. Our current identification of *elavl4, rorb, cbpc6*, and *epn4* is primarily based on genomic proximity and bioinformatic inference. The specific mechanistic roles of these genes in host-pathogen interactions during *Vibriosis* have yet to be experimentally confirmed. Future research should leverage targeted approaches such as CRISPR/Cas9 gene editing, RNA interference (RNAi), or heterologous overexpression in cell lines and *in vivo* models. Such functional characterizations have already been successfully executed in the yellow drum for candidates including *NK-lysin* (Huang et al., 2023), *Galectin-9* (Luo et al., 2023), and *Cygb* (Zhou et al., 2023), providing a robust methodological template for our ongoing work. Third, the phenotypic assessment in this study relied on a single, controlled artificial challenge. While such experiments are standardized and highly reproducible, they may not fully encapsulate the environmental complexity and multi-pathogen dynamics of natural infections in commercial aquaculture settings (Hu et al., 2020). Future studies would benefit from incorporating longitudinal data from natural outbreaks or multiple challenge trials across different life stages to refine the accuracy of phenotypic evaluations. Finally, the translation of these genomic findings into breeding programs is the ultimate objective. The significant SNP and InDel markers identified herein must be converted into cost-effective high-throughput genotyping tools (e.g., KASP or targeted sequencing panels). Their predictive accuracy needs to be rigorously evaluated in diverse breeding kernels. Integrating these high-confidence markers into Genomic Selection (GS) models could significantly accelerate the development of *V. harveyi*-resistant lines in *P. diacanthus*. GS has already demonstrated substantial efficiency gains over traditional pedigree-based methods in various aquaculture species, such as the Pacific oyster (Yang et al., 2024) and common carp (Palaiokostas et al., 2019), underscoring the transformative potential of genomic-informed breeding in the sciaenid industry.

## Conclusion

In conclusion, this study represents the first integrated genome-wide association study (GWAS) and genetic differentiation analysis (*F*_ST_) to systematically dissect the genetic basis of *V. harveyi* resistance in the blackspotted croaker (*Protonibea diacanthus*). Our findings demonstrate that resistance to vibriosis is a complex trait under moderate-level genetic control, with genomic heritability estimated at 0.374 ± 0.29 for survival time (ST) and 0.237 ± 0.185 for survival status (SS). The near-perfect genetic correlation identified between ST and SS (*r*_*g*_ = 0.999) suggests that these two metrics are governed by an almost identical genetic architecture, while the negligible correlation with body weight indicates that disease resistance and growth performance are genetically independent. Through the convergence of GWAS and selection sweep signals, we identified a pivotal QTL on chromosome 4 and prioritized a set of high-confidence candidate genes, including *elavl4, rorb, cbpc6*, and *epn4*. Functional enrichment analysis further revealed that resistance in *P. diacanthus* is significantly associated with RNA metabolic processes, mRNA surveillance, and nucleocytoplasmic transport, highlighting the crucial role of post-transcriptional regulation in host-pathogen interactions. These results not only fill a major knowledge gap in the genomics of *P. diacanthus* but also provide essential molecular targets for marker-assisted selection (MAS) and genomic selection (GS). Ultimately, this research provides a robust genomic framework for the development of disease-resistant strains, facilitating the sustainable development and high-quality expansion of the blackspotted croaker aquaculture industry.

## Author contribution

WLC: conceptualization, supervision, funding acquisition, and review and editing. SS: sample collection, data analysis, and manuscript writing. LZ: sample collection and data analysis. NL, PX: project coordination and manuscript revision. LY, KQY: sample collection, provision of breeding sites and experimental fish. The authors declare no competing interests.

