## Supplementary material for "Genome-wide association study and genetic differentiation analysis reveal the genetic basis of *Vibrio harveyi* resistance in blackspotted croaker (*Protonibea diacanthus*)": Figure S1A, Figure S1B

**Supplementary materials**

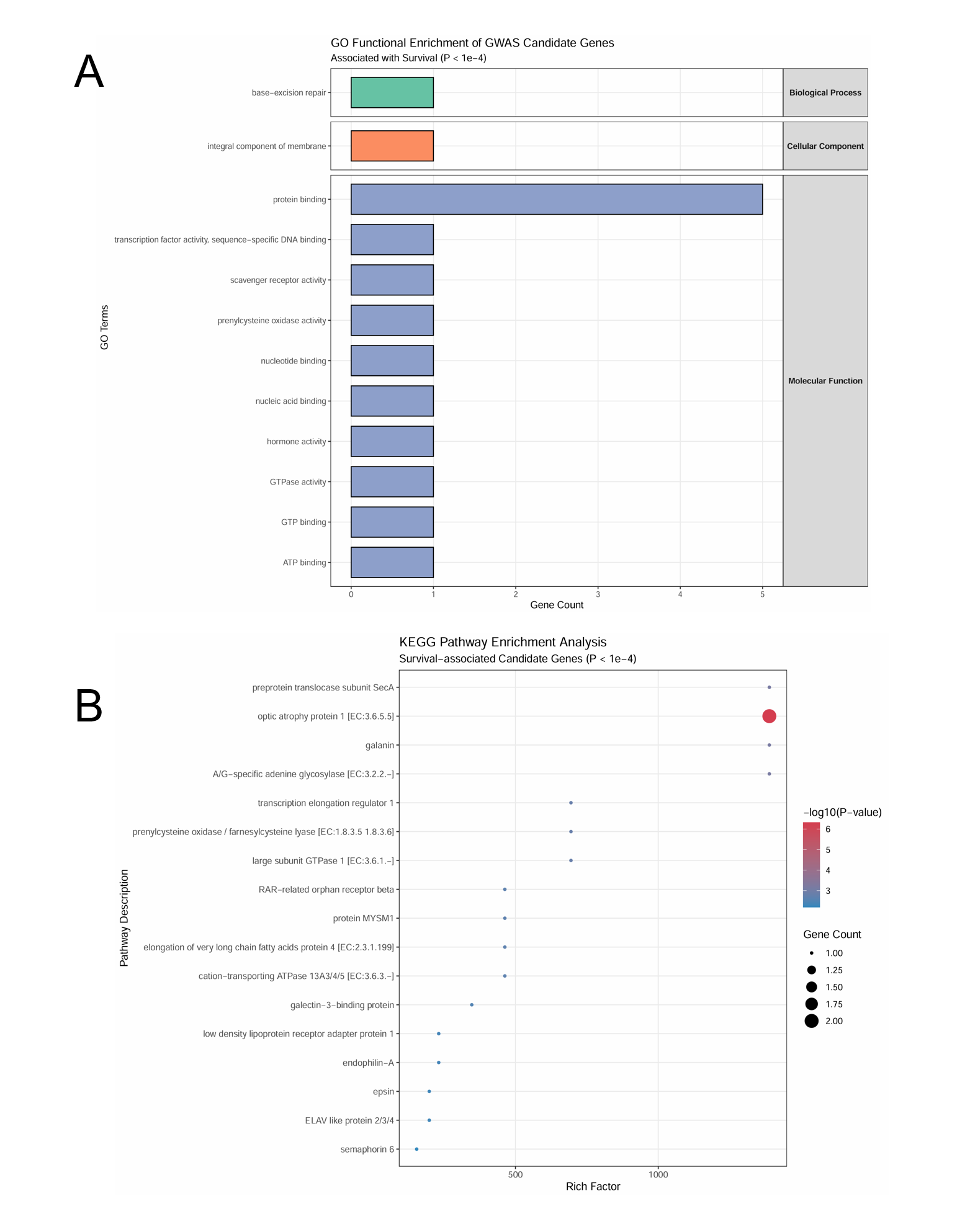

FIGURE S1 Enriched GO and KEGG terms for the candidate genes of *V. harveyi* resistance. A: GO enrichment. B: KEGG enrichment.
